# Palmitoleate Protects against Zika virus infection-induced Endoplasmic Reticulum Stress and Apoptosis in Neurons

**DOI:** 10.1101/2025.01.22.634157

**Authors:** Chandan Krishnamoorthy, Anthony Delaney, Devanshi Shukla, Taija Hahka, Ann Anderson-Berry, Sathish Kumar Natarajan

## Abstract

Zika virus (ZIKV) infection during pregnancy is associated with the development of fetal complications such as microcephaly. We have recently demonstrated that palmitoleate protects against ZIKV-induced apoptosis in placental trophoblasts. In the present study, we hypothesize that palmitoleate prevents ZIKV infection-induced endoplasmic reticulum (ER) stress and apoptosis in neurons. Neurons were infected with 0.1-1 multiplicity of infection of recombinant MR766 or PRVABC59 strains of ZIKV for an hour followed by treatment of palmitoleate (100 µM-200 µM) for different post-infection time points. Apoptosis was measured by nuclear morphological changes, caspase 3/7 activity, and immunoblot analysis of pro-apoptotic mediators. Activation of ER stress markers and viral envelope levels were detected using qRT-PCR and immunoblot analysis. Infectious virus particles were measured by using plaque assay. ZIKV infection to neuronal cells showed increased levels of pro-apoptotic markers like cleaved-PARP, cleaved caspase-3, Bim, and Puma, whereas decreased levels of anti-apoptotic markers such as Mcl-1, Bcl-1, and Bcl-xL. Further, we observed activation of three arms of ER stress namely: inositol requiring enzyme 1 alpha (IRE1), protein kinase-like ER kinase (PERK), and activating transcription factor (ATF6) pathways with ZIKV infection. Treatment of palmitoleate dramatically decreased ZIKV infection-induced increase in percent apoptotic nuclei and caspase 3/7 activity. Further, treatment of palmitoleate decreased cleaved PARP and PUMA protein expressions. Treatment of palmitoleate reduced ZIKV-induced ER stress activation as evidenced by decreased levels of phosphorylated forms of IRE1 and eukaryotic initiation factor 2 alpha; decreased expressions of cleaved ATF6, spliced X-box associated protein 1 and C/EBP homologous protein compared to ZIKV infection alone. Further, treatment of palmitoleate attenuated ZIKV envelope levels and infectious titer in SH-SY5Y and primary fetal cortical neurons isolated from humanized STAT2 knockin mice. These data suggest that palmitoleate supplementation protects against ZIKV-induced neuronal ER stress, apoptosis and decreases Zika viral load thereby mitigates neuronal damage.

## INTRODUCTION

Zika virus (ZIKV) belongs to the *Flaviviridae* family, genus *Flavivirus,* and is an enveloped, single-stranded positive-sense RNA virus(1). ZIKV is approximately 50 nm in diameter with structural and non-structural proteins(2). Three structural proteins, namely envelope, capsid, and membrane proteins play an important role in the formation of viral particles. Seven non-structural (NS) proteins, namely NS1, NS2A, NS2B, NS3, NS4A, NS4B, and NS5 are involved in the viral replication, processing, and viral assembly(1, 3).

ZIKV can spread to humans primarily from *Aedes* genus mosquito bites; especially *Aedes aegypti*(4). It can also be transmitted through sexual contact, during pregnancy from the maternal circulation to the fetus, *in utero*, and through the ZIKV present in the breast milk(3, 5). Clinical symptoms of ZIKV include headache, rashes, mild fever, edema, and retro-orbital pain accounting for 20% of infected populations, whereas 80% are asymptomatic(6). Women in their first and second trimesters of pregnancy are more prone to Zika viral infections(3). The placenta develops during the first trimester allowing proper nutrient and oxygen transfer from mother to fetus. ZIKV infection to the placental cells further multiplies and increases its ability to reach and spread to the fetus, especially affecting the neuronal cells in the fetal brain(3).

Pregnancy-related vertical transmission of ZIKV from mother to fetus results in the development of Congenital Zika Syndrome(7). The key symptoms of congenital Zika syndrome are microcephaly, contractures, hypertonia, calcification of the subcortex, thinning of the cortex, and retinal pigmentary mottling(7). A large-scale cohort study from Brazil during the 2016 ZIKV epidemic shows that congenital Zika syndrome in children with maternal ZIKV infection had a higher risk for death compared to children without congenital Zika syndrome(8). Pathogenesis of ZIKV has revealed that its infection during the early stage of fetal development results in extensive apoptosis of neuronal progenitor cells(9). ZIKV infection induces activation of extrinsic and intrinsic pathways of apoptosis and endoplasmic reticulum (ER) stress-mediated apoptosis(10–12).

The ER has multiple functions in a cell such as protein synthesis, secretion, and folding. ER is also involved in the biosynthesis of lipids, and steroids and helps in maintaining calcium homeostasis. ZIKV is dependent on the host cellular machinery for its replication in the ER and ER-related proteins are shown to be essential for viral replication(13). As a result, ZIKV infection leads to excessive misfolding and unfolding of protein accumulation in the ER, thereby triggering ER stress. Sustained activation of the three arms of the ER transmembrane proteins can activate a cascade of downstream pro-apoptotic signaling such as C/EBP homologous protein (CHOP) or c-Jun N-terminal kinase (JNK) for the initiation of apoptosis(14).

Many viral infections during pregnancy can cause severe fetal abnormalities; however safe and therapeutic antivirals are scarce. Specifically, there are no FDA-approved vaccines or antivirals available for the ZIKV infection. An alternative approach is the use of dietary nutrient compounds that can be easily accessible, affordable, and safe to administer during pregnancy. Specifically, we used palmitoleate (PO), an omega-7 monounsaturated fatty acid to mitigate ZIKV infection. Our previous studies have shown that ZIKV infection induces sustained ER stress- and mitogen-activated protein kinase (MAPK) activation-dependent apoptosis in placental trophoblast(12). Treatment of PO shows protective effects against ZIKV-induced ER stress, MAPK, and apoptosis in trophoblasts(15, 16). In the present study, we hypothesized that ZIKV infection would induce ER stress and apoptosis in neuronal cells, and we have also tested the protective role of PO against ZIKV-induced ER stress and apoptosis in neuronal cells.

## MATERIALS AND METHODS

### Materials

Chemicals and buffers were procured from ThermoFisher Scientific (Massachusetts, USA). Palmitoleate (# P9417) and Bovine Serum Albumin ( # A3803) were obtained from Sigma-Aldrich, MO, USA. Pan caspase inhibitor Z-VAD-FMK (# S7023), IRE1α endonuclease inhibitor (# STF-083010), and Salubrinal (eIF2α dephosphorylation inhibitor) (# S2923) were acquired from Selleckchem, TX, USA. Apo-ONE® Homogeneous Caspase-3/7 Assay (# G7792), QIAamp viral-RNA extraction kit (# 52906), LightCycler® 480 SYBR Green I Master (# 04707516001), and Nitrocellulose membrane, 0.2 µm (# 1620112) were obtained from Promega, Qiagen, Roche, and Bio-Rad, respectively. Novex™ Tris-Glycine Mini Protein Gels (# XP00100BOX, # XP00102BOX, # XP00120BOX) from Invitrogen, CA, USA.

### Antibodies

Primary antibodies for PARP (# 9542), Cleaved Caspase-3 (# 9661), Phospho-eIF2α (# 3398S), eIF2α (# 5324S), IRE1α (# 3294S), Bim (# 2933S), Bcl-2 (# 15071S), Bcl-xL (# 2764S), Mcl-1 (# 94296S), and XBP-1s (# 40435S) were acquired from Cell Signaling Technologies, MA, USA. Anti-IRE1 (phospho S724) (#ab48187), and Anti-ATF6 antibody (# ab122897) were purchased from Abcam, Cambridge, UK. PUMAα/β Antibody (# sc-28226), monoclonal anti-β-Actin antibody (# A5441), and Zika virus envelope protein antibody (# GTX133314) were obtained from Santa Cruz Biotechnology (TX, USA), MilliporeSigma (MA, USA), and Genetex (CA, USA), respectively. Secondary antibodies such as peroxidase affiniPure™ Goat Anti-Rabbit IgG (H+L) and Donkey Anti-Mouse IgG (H+L) were obtained from Jackson Immuno Research (PA, USA).

### Cell culture

Cell lines were obtained from ATCC. SH-SY5Y (# CRL-2266), neuroblastoma-derived neuronal-like cells (referred to as neuronal cells in this manuscript) were cultured in MEM (# 10-024-CV, Corning, NY, USA) and F12 (# 11-765-047, Gibco) in a 1:1 ratio with 10% FBS (# 10437-028, Gibco) and 0.01% plasmocin (# MPT-43-02, CA, USA). The media was changed every 2-3 days. Vero cell line (# CCL-81) was cultured in DMEM (# 10-013-CV, Corning, NY, USA), 10% FBS, and 0.01% plasmocin. Cell lines were maintained at 37^◦^C with 5% CO2 and 95% relative humidity.

### Humanized STAT2 knockin (hSTAT2KI) mice model

Humanized STAT2 knockin (hSTAT2KI) mice were obtained from Dr. Michael S. Diamond, Washington University of School of Medicine, St Louis, MO, USA. The protocol was approved by the Institutional Animal Care and Use Committee (IACUC) at the University of Nebraska-Lincoln (UNL), and the animals were housed in the Life Science Annex at the UNL. Eight-week-old female and male hSTAT2KI mice were mated and at gestational day 13.5 female mice were sacrificed and cortical neurons were isolated from the fetal brain.

### Isolation of primary cortical neurons from hSTAT2KI mice fetal brain

Mice fetal cortical neurons were isolated according to Wu and Gorantla, 2013(17). Briefly, the cortical tissue was isolated from hSTAT2KI mice fetal brains. D-PBS (# SH30028.02, Cytiva), 0.25% trypsin (# 25-053-CI, Corning, NY, USA), and DNase (# D5025-375KU, Sigma-Aldrich, MO, USA) were used for cell dissociation and cultured in a neurobasal medium (# 21-103-049, Gibco) supplemented with B-27 serum-free supplement (# 17-504-044, Gibco), GlutaMAX (# 35-050-061, Gibco), 100 I.U./mL penicillin and 100 μg/mL streptomycin (# 15140122, Gibco). The media was replaced every two days and on day 12 the neurons were used for the experiment.

### ZIKV Infections

ZIKV strains namely, recombinant clone of MR766 (rMR) and PRVABC59 (PR) were obtained from Dr. Pattnaik’s lab, University of Nebraska-Lincoln, NE, USA, and used for infection at MOI of 0.1 to 1.0 for an hour using viral infection media. The viral infection media contain appropriate culture media for each cell type (SH-SY5Y: MEM and F12; hSTAT2KI mice cortical neuron: Neurobasal medium, B-27 serum-free supplement, and GlutaMAX) with 2% FBS, 1× MEM Non-Essential Amino Acids Solution (# 11140050, Gibco), 2mM Glutamine (# 25030149, Gibco), 10 mM HEPES (# H0887, Sigma-Aldrich), 100 I.U./mL penicillin and 100 μg/mL streptomycin. After infection, the media was replaced with 10% FBS-containing growth media, and the infected cells were analyzed after post-infection time points.

### Treatment of Palmitoleate (PO)

A palmitoleate stock concentration of 80 mM was prepared using isopropanol. BSA (1%) was prepared by dissolving in growth media and incubated at 37^◦^C for 30 minutes to completely dissolve the BSA and filter sterilization using the sterile filter. Palmitoleate (PO) with 100 μM and 200 μM final concentrations was prepared using the 1% BSA-containing media and incubated at 37^◦^C for 20 minutes for the fatty acid-BSA conjugation. Later, ZIKV-infected cells were treated with PO and analyzed after post-infection time points.

### Characterization of apoptosis

Biochemical and structural markers of apoptosis were analyzed by caspase 3/7 activity and percent apoptotic nuclei, respectively. Cells were seeded in 24 well plates, after ZIKV infection, and at different time points of treatment, cells were stained with 5 μg/ml of 4’, 6-diamidine-2-phenylindole dihydrochloride (DAPI, # D9542, Sigma-Aldrich, MO, USA) for 10 minutes at 37^◦^C, and the nuclear morphology changes (condensation and fragmentation) were observed using an epi-fluorescence microscope, and percent apoptotic nuclei were calculated. The experiments were repeated in triplicate and at least 100 cells per well. Caspase 3/7 activity was calculated using Z-DEVD-R110 [Rhodamine 110 bis-(N-CBZ-I-aspartyl-I-glutamyl-I-Valyl-aspartic acid amide] as a substrate. Z-DEVD-R110 can be cleaved by both caspase 3 and 7 enzyme activities in the cells which releases rhodamine 110 fluorophore and is measured at 498 nm excitation and 521 nm emission (BioTek Synergy). Experiments were performed in quadruplicate and caspase 3/7 activity is calculated according to the manufacturer’s instructions. The results were represented as the net fluorescence fold changes compared to the vehicle-treated cells.

### RNA isolation and Quantitative Polymerase Chain Reaction (qPCR)

Total RNA was extracted from the cells using TRIzol (# 15596018, Invitrogen, MA, USA) and a QIAamp viral-RNA extraction kit was used to isolate the viral RNA from the cell culture supernatant. cDNA was synthesized with 0.5 - 1 µg of RNA using random primer mix (# S1330S, NEB, MA, USA), Deoxynucleotide (dNTP) Solution Mix (# N0447L, NEB, MA, USA), M-MuLV Reverse Transcriptase Reaction Buffer (# B0253S, NEB, MA, USA), 0.1 M DTT (# Y00147, Invitrogen, MA, USA), RNase Inhibitor (# M0307L, NEB, MA, USA) and M-MuLV Reverse Transcriptase (# M0253L, NEB, MA, USA). To quantify viral envelope RNA copy number the primers targeting the E gene of ZIKV were used with the hydrolysis probe. The absolute expression of envelope RNA copy numbers was quantified using a standard curve from a custom-synthesized ZIKV PCR product. mRNA expression of CHOP, s-XBP1, us-XBP1, ATF4, and BiP levels in cells was quantified relative to 18S rRNA using LightCycler® 480 SYBR Green I Master in a CFX Connect Real-Time PCR Detection System (Bio-Rad). The list of primers and probe used are listed in Table 1.

**Table 1.**
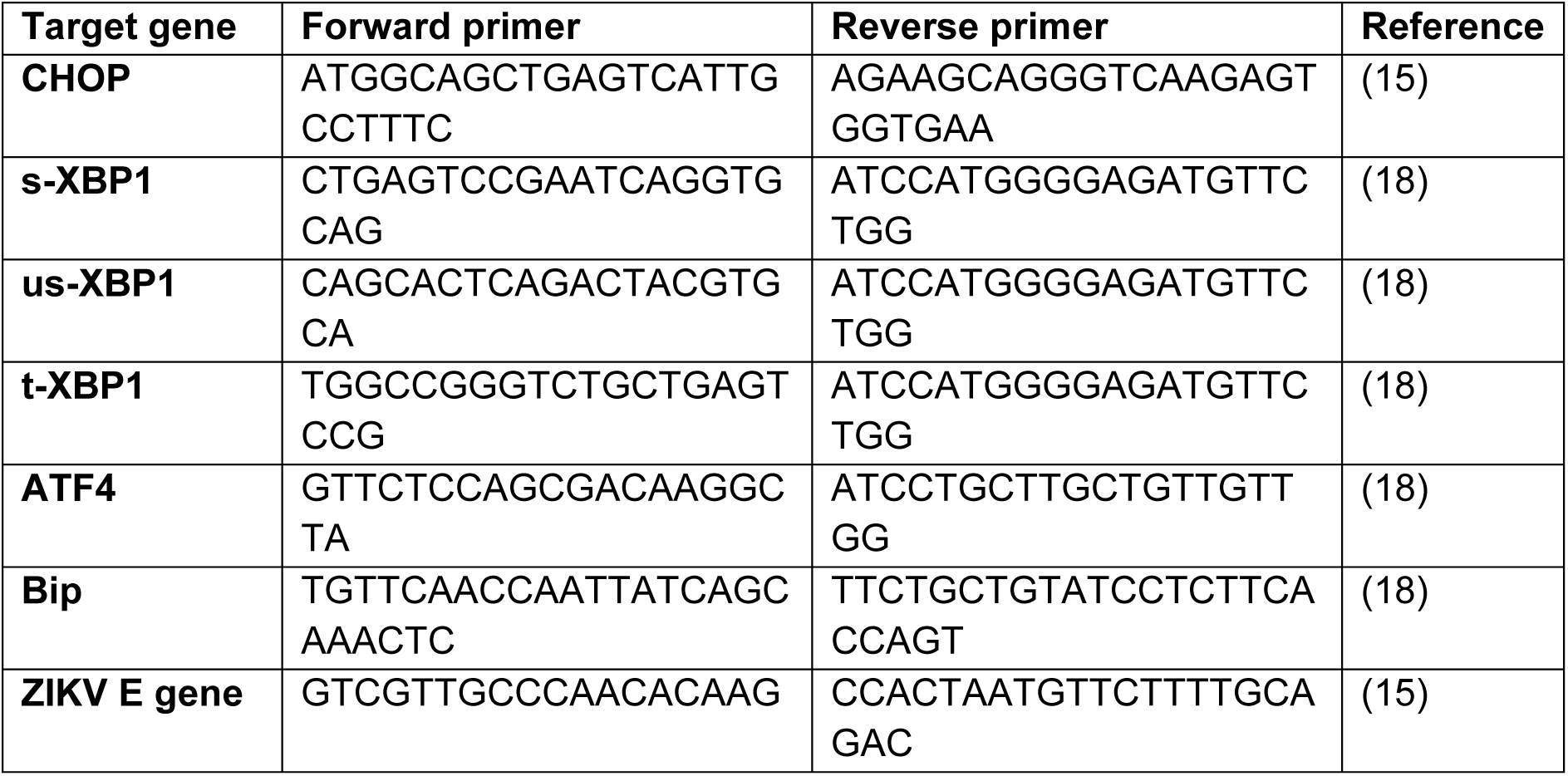
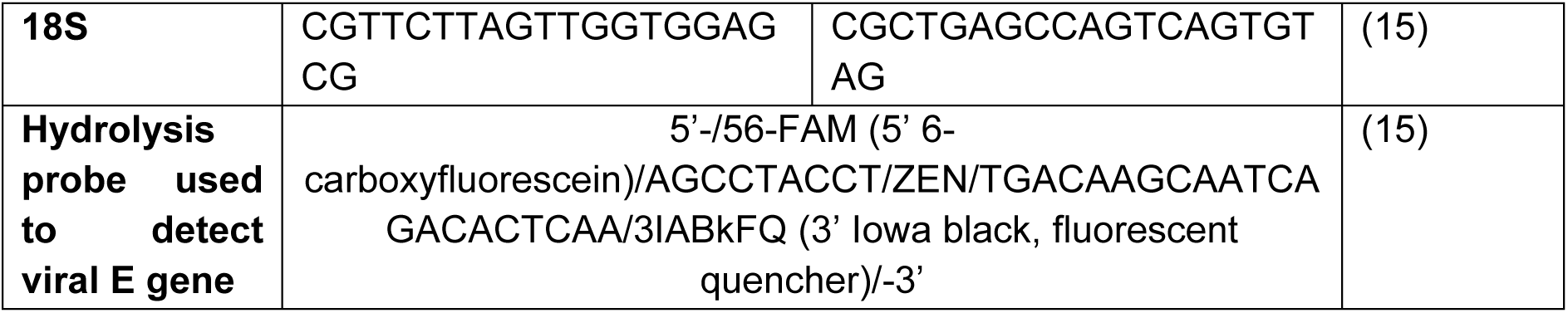
List of primers and probe.

### Immunoblot analysis

The cell culture media was removed, washed with ice-cold 1x PBS and the lysis buffer (100 µl) was added to the cells followed by cell scraping on ice. The lysis buffer was prepared using NaCl (150 mM), DTT (1 mM), Tris (50 mM) at pH 7.4, Na_3_VO_4_ (1 mM), EDTA (1 mM), NaF (100 mM), PMSF (1 mM), Triton x-100 (1%), protease and phosphatase inhibitor cocktail (# 78440, Thermo Scientific, MA, USA). The cell lysate was centrifuged at 15,000 × g for 20 min and the supernatant was used for the protein estimation by Pierce 660-nm Protein assay reagent (# 22660, Thermo Scientific, MA, USA). Around 15 to 30 µg of total proteins were separated using SDS-PAGE and transferred to the nitrocellulose membrane using a transfer buffer at 4^◦^C. BSA (5%) (# BP1600-100, Fisher Scientific, NH, USA) in TBST was used for blocking and diluting the primary (1:1000) and secondary (1:5000) antibodies. Blocking was done for 1 h at 37^◦^C, primary antibody incubation was overnight at 4^◦^C, and secondary antibody incubation was at 37^◦^C for 2 h at RT. The blots were washed with TBST 3 times, at 10-minute intervals after the primary and secondary antibody incubations. Western Lightning Plus, Chemiluminescent Substrate (# NEL104001EA, Revvity Health Science Inc.) was used to develop the immunoblots and imaged using ChemiDoc Touch Imaging System (Bio-Rad).

### Immunofluorescence staining and imaging

SH-SY5Y cells were infected with 1 MOI rMR for 48 h of post-infection. Briefly, the media was removed, and cells were washed with 1X PBS, fixed for 30 minutes at 37◦C using 3% paraformaldehyde in PBS with 100 mM PIPES, 3 mM MgSO4, 1 mM EGTA, and washed with 1X PBS thrice. Cell permeabilization was done with 0.3% Tween 20 in PBS for 15 minutes at room temperature and washed with 1X PBS thrice. Cells were blocked with 5% glycerol, 5% goat serum, and 0.01% sodium azide in PBS for 60 minutes at 37◦C and washed with 1X PBS thrice. Primary and secondary antibodies were prepared using a blocking buffer. Primary antibody incubation was done in 1:200 dilution at 4°C overnight and washed three times with 1X PBS, whereas the secondary antibody (Alex conjugated) was for 1 hour at 37◦C. Then the cells were washed with 1X PBS and deionized water once and counterstained with DAPI. Finally, the cells were washed with PBS twice and the Fluoromount G (# 17984-25, Electron Microscopy Sciences, PA, USA) was used to mount on the microscope slides. Images were acquired using a Nikon A1R-Ti2 confocal system.

### Plaque assay

Infectious viral titer quantification from hSTAT2KI mouse cortical neuron and SH-SY5Y was performed by Plaque assay(19). Infection media was prepared using DMEM (1X), FBS (2%), NaHCO3 (0.0023%), HEPES (2%), Sodium pyruvate (1%), Pen/Strep (1%), and non-essential amino acids (1%). The cell culture supernatants were diluted 10 folds using the infection media, 10^-3^ and 10^-4^ dilutions were used in triplicate. The media was removed from the 90% confluent Vero cells, washed with 1X PBS, and the diluted samples were plated for the virus adsorption for an hour in 12 well plates with gentle rocking every 5 minutes. After adsorption, the infection media was removed and the cell monolayer was overlaid with medium prepared by mixing 2% low melting agarose (# BP165-25, Fisher Scientific, NH, USA) with equal volume of media containing 2 X DMEM (# SLM-202-B, Sigma-Aldrich, MO, USA), FBS (4%), NaHCO3 (0.0046%), HEPES (4%), Sodium pyruvate (2%), Pen/Strep (2%), non-essential amino acids (2%), and Plasmocin (0.02%). The plates were incubated at 37°C for 5 days followed by fixation for 30 minutes using 10% neutral buffered formaldehyde and the agarose plugs were removed and stained for 10 minutes using 0.1% crystal violet dissolved in 30% methanol. The plates were washed with distilled water and allowed to air dry. Plaques were counted manually, and the results were expressed in Plaque Forming Units/mL (PFU/mL).

### Quantification and statistical analysis

The statistical analysis was performed using GraphPad Prism version 9.5.1. The data were analyzed using a one-way analysis of variance with multiple comparisons with Bonferroni post hoc corrections and the data were represented as mean ± SEM. Immuno blots were analyzed and quantified using Image Lab and ImageJ software, respectively.

## RESULTS

### ZIKV induced caspase-dependent apoptosis with an increase in pro-apoptotic and a decrease in the anti-apoptotic mediators in neuronal cells

#### ZIKV induces caspase-dependent apoptosis

Neuronal cells were infected with a recombinant clone of African strain (rMR) or Asian strain (PRV) of ZIKV with 0.1, 0.25, 0.5, 0.75, and 1.0 multiplicity of infections (MOIs), and at 96 h of post-infection, structural and biochemical markers of apoptosis such as percent apoptotic nuclei and caspase 3/7 activity, respectively were measured. We observed an increase in percent apoptotic nuclei and caspase 3/7 activity with an increasing MOI in both rMR and PR strain-infected neuronal cells. rMR infection showed a significant increase in percent apoptotic nuclei starting from 0.5 – 1.0 MOI, whereas a significant increase in percent apoptotic nuclei with PR infection was only observed at 1.0 MOI after 96 h of post-infection (**Fig. 1A, B**). Both r-MR and PR show a significant increase in caspase 3/7 activity from 0.25 - 1.0 MOI compared to mock-infected neuronal cells. (**Fig. 1A, B**). These data indicate that ZIKV induces apoptosis and that rMR exhibited greater virulence than the PR strain in inducing apoptosis to neuronal cells at lower MOIs.

**Fig. 1.**
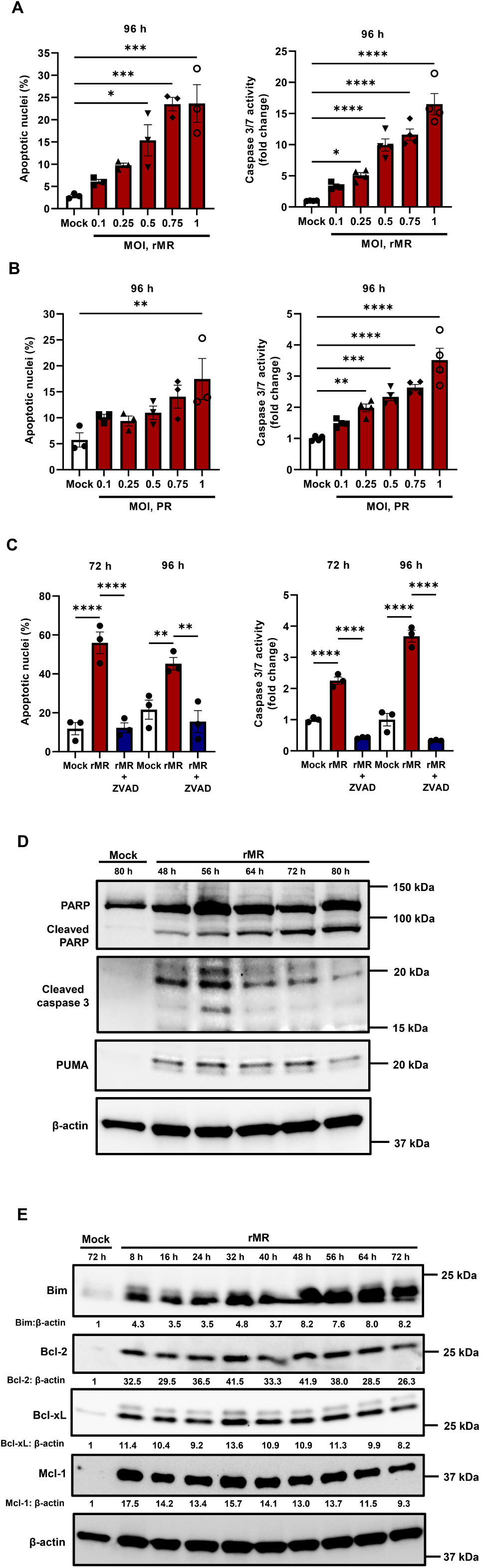
ZIKV induced caspase-dependent apoptosis with an increase in pro-apoptotic and a decrease in the anti-apoptotic mediators in neuronal cells. **A, B.** SH-SY5Y cells were infected with recombinant MR766 (rMR) and PRVABC59 (PR),0.1 - 1 MOI up to 96 hours of post-infection (hpi). For both ZIKV strains increase in MOI increased the present apoptotic nuclei and caspase 3/7 activity compared to the mock. **C.** Treatment of pan-caspase inhibitor, Z-VAD-FMK, (Z-VAD) significantly decreased the percent apoptotic nuclei and caspase 3/7 activity in 1 MOI rMR infected cells at 72 and 96 h post-infection time points. **D.** Immunoblot analysis showed cells infected with 0.1 MOI rMR had a drastic increase in the cleaved PARP at 48 h - 80 h and cleaved caspase 3, PUMA at 48 h - 72 h of post infections. **E.** Cells infected with 0.1 MOI rMR at different post-infection time points (8 h - 72 h) showed a time-dependent increase in the Bim expression, and a decrease in expressions of anti-apoptotic markers (Bcl-2, Bcl-xL, Mcl-1). Beta-actin was used as loading control and remained unchanged. The images are representative images. Data presented as mean ± SEM, n=3 or 4 *p<0.05, ** p<0.01, *** p<0.001, **** p<0.0001 compared to mock or ZIKV infected cells

To determine whether ZIKV induces caspase-dependent neuronal apoptosis, we treated ZIKV-infected cells with a pan-caspase inhibitor, Z-VAD-FMK. At 72 h and 96 h of post-infection time points, rMR infection of 1.0 MOI showed a significant increase in percent apoptotic nuclei and caspase 3/7 activity compared to the mock infections and treatment of Z-VAD-FMK significantly attenuates ZIKV-induced neuronal apoptosis (**Fig. 1C**). To further confirm the role of caspase activation with ZIKV infection, we measured the levels of caspase substrates such as cleaved PARP and cleaved caspase 3 by immunoblot analysis in the cells infected with 0.1 MOI rMR, 48 - 80 h post-infection. As post-infection time increased, levels of cleaved PARP were also increased. The expression of cleaved PARP was remarkably higher compared to mock infection, specifically after 64-80 hours post-infection (hpi) (**Fig. 1D**). Increased cleaved caspase 3 at 19 kDa was observed at all time points tested (48-80 hpi) with ZIKV infection (**Fig. 1D**). The highest abundance of 19 kDa, cleaved caspase 3 was observed between 48 h and 56 hpi, and their levels slightly decreased in 64 - 80 h of post-infection compared to 48-56 h time points. We also observed an increased level of 17 kDa band of cleaved caspase 3 at 56 hpi (**Fig. 1D**). Further, increased levels of PUMA, a pro-apoptotic BH3 domain-containing protein, were observed between 48 - 72 hpi time points compared to the mock infection and the levels of PUMA decreased by 80 h (**Fig. 1D**). These data demonstrate that ZIKV infection induces caspase-dependent apoptosis in neuronal cells.

#### ZIKV infection alters pro-apoptotic and anti-apoptotic mediators

To determine the role of pro-apoptotic and anti-apoptotic mediators with ZIKV infection in neuronal cells, we infected SH-SY5Y cells with 0.1 MOI rMR and measured the levels of Bim, Bcl-2, Bcl-xL, and Mcl-1. Pro-apoptotic protein marker, Bim expression was highly increased from 48 - 72 h compared to mock and earlier time points tested (8 - 40 h) (**Fig. 1E**). However, anti-apoptotic and BH-3 domain-containing proteins such as Bcl-2, Bcl-xL, and Mcl-1 levels were highly increased in ZIKV infected neuronal cells at the initial time point 8 - 56 hpi, compared to the later time point 64 - 72 hpi (**Fig. 1E**). These data indicate that increased levels of pro-apoptotic and decreased expression of anti-apoptotic markers were observed in ZIKV-infected neuronal cells after 64 - 72 hpi.

### ZIKV infection induces sustained ER stress in neuronal cells

To determine the role and degree of activation of ER stress with ZIKV infection, we analyzed key markers of ER stress such as phosphorylation of IRE1α, eIF2α, ATF6 cleavage, and XBP1 splicing. ZIKV (rMR) infected cells showed increased levels of p-IRE1α at 8 h to 32 h of post-infection time points compared to the mock infection (**Fig. 2A**). We also observed an increase in the full-length and cleaved ATF6 levels at 8 - 32 hpi (**Fig. 2A**). The activation of downstream effectors for PERK and IRE1α was detected by measuring levels of phosphorylated eIF2α (p-eIF2α) and spliced-XBP1 (s-XBP1), respectively. ZIKV-infected cells at 72 h of post-infection showed an increase in p-eIF2α and s-XBP1 expression compared to the mock infection (**Fig. 2B**) and at 96 h showed a dramatic increase in p-eIF2α and s-XBP1 protein expression (**Fig. 2B**). These data suggest that ZIKV infection induces activation of ER stress in the neuronal cells.

**Fig. 2.**
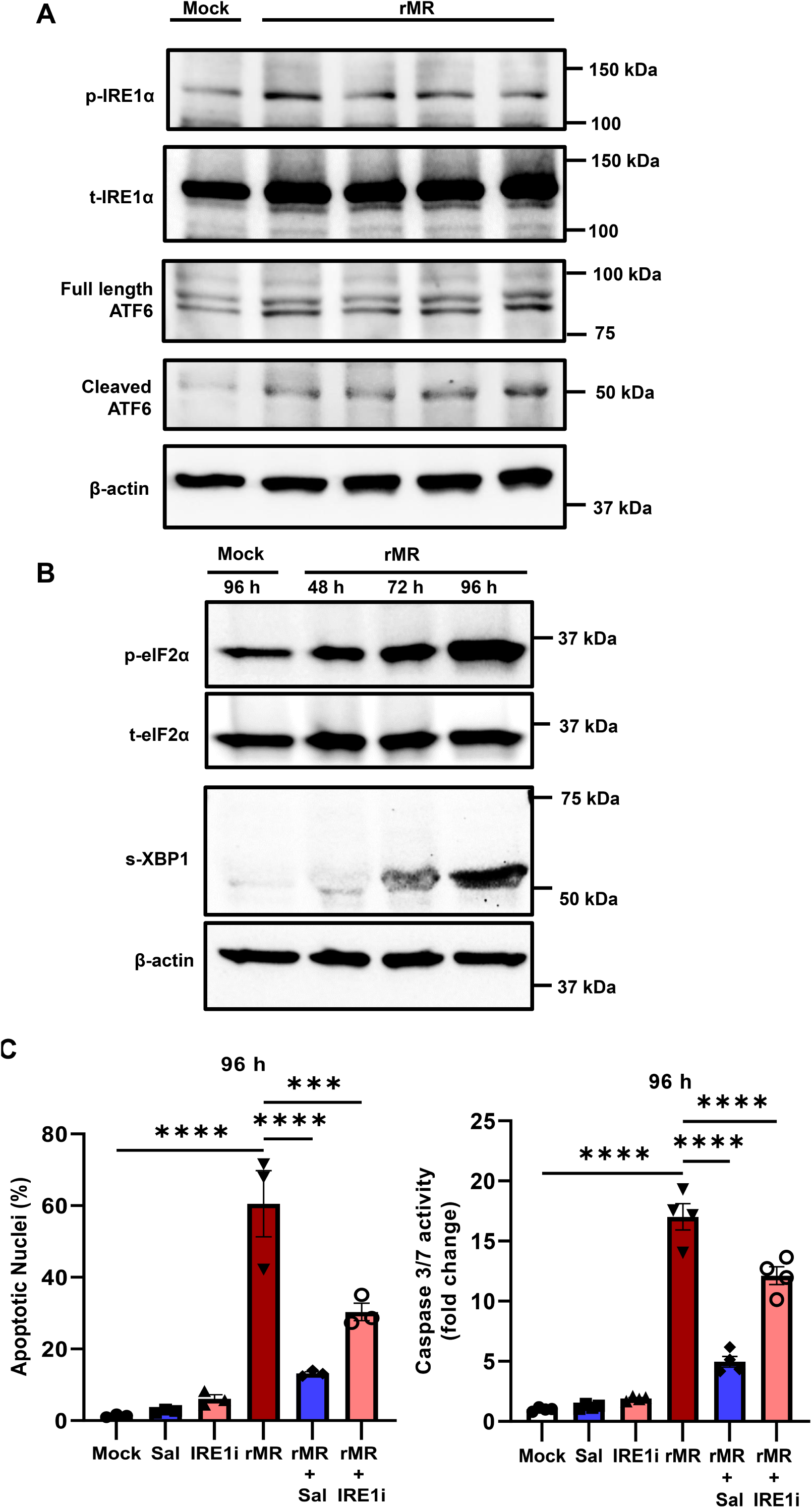
ZIKV infection induces sustained ER stress in neuronal cells. **A.** Immunoblot analysis showed an increase in p-IRE1α, increase in full length, and cleaved ATF6 at 8h – 32 h of post-infection time point when infected with 0.1 MOI rMR. **B.** An increase in p-eIF2α and s-XBP1 was detected at 72 h and 96 h of post-infections when the infected with 0.1 MOI rMR. Total (t)IRE1α and t-eIF2α levels were unaltered between mock and rMR-infected neuronal cells. **C.** Cells treated with Salubrinal (eIF2α dephosphorylation inhibitor, eIF2αi), and STF-083010 (IRE1αi, IRE1α endonuclease inhibitor) significantly decreased the apoptotic nuclei percentage and caspase 3/7 activity after 96 h of post-infection. Beta-actin was used as loading control and remained unchanged. The images are representative images. Data presented as mean ± SEM, n=3 or 4, *** p<0.001, **** p<0.0001 compared to mock or ZIKV infected cells.

To evaluate the critical role of ER stress in the ZIKV-induced apoptosis, we used small molecule inhibitors namely Salubrinal (eIF2α dephosphorylation inhibitor, eIF2αi), and STF-083010 (IRE1α endonuclease inhibitor, IRE1αi). Neuronal cells were infected with 1 MOI rMR at 96 h of post-infection with and without small molecule inhibitors of ER stress and apoptosis was determined. ZIKV infection at 96 h showed significantly increased percent apoptotic nuclei and caspase 3/7 activity (**Fig. 2C**). Neuronal cells treated with eIF2α inhibitor and IRE1α inhibitor had significantly reduced percent apoptotic nuclei and caspase 3/7 activity (**Fig. 2C**). These data disclose the critical role of ER stress activation and inhibition of eIF2α and IRE1α significantly attenuates ZIKV-induced apoptosis in neuronal cells.

### Palmitoleate (PO) protects against ZIKV-induced apoptosis in neuronal cells

To determine the protective role of palmitoleate (PO), we infected neuronal cells with 1 MOI of rMR or PR followed by treatment of PO (100-200 µM) for 96 h, and apoptosis was measured. Both rMR and PR strain infection with 1 MOI showed a significant increase in percent apoptotic nuclei and caspase 3/7 activity (**Fig. 3 A, B**). Interestingly, treatment of PO significantly reduced the percent apoptotic nuclei and caspase 3/7 activity in both rMR and PR infection (**Fig. 3 A, B**). The protective role of PO was analyzed by measuring the levels of cleaved PARP and PUMA expression in ZIKV-infected neuronal cells. At 72 h of post-infection time point, cells infected with 0.1 MOI rMR exhibited an increase in cleaved PARP and PUMA, whereas treatment with PO (100 µM and 200 µM) decreases the levels of these proteins (**Fig. 3C**). Further, at 96 h post-infection, a dramatic increase in cleaved PARP was observed with 0.1 MOI rMR and treatment of PO (100 µM and 200 µM) considerably decreased the expression of cleaved PARP (**Fig. 3D**). PR infection (0.1 MOI) also increased the levels of cleaved PARP and treatment of 200 µM of PO drastically decreased the expression (**Fig. 3E**). Together, palmitoleate prevents ZIKV-induced neuronal apoptosis.

**Fig. 3.**
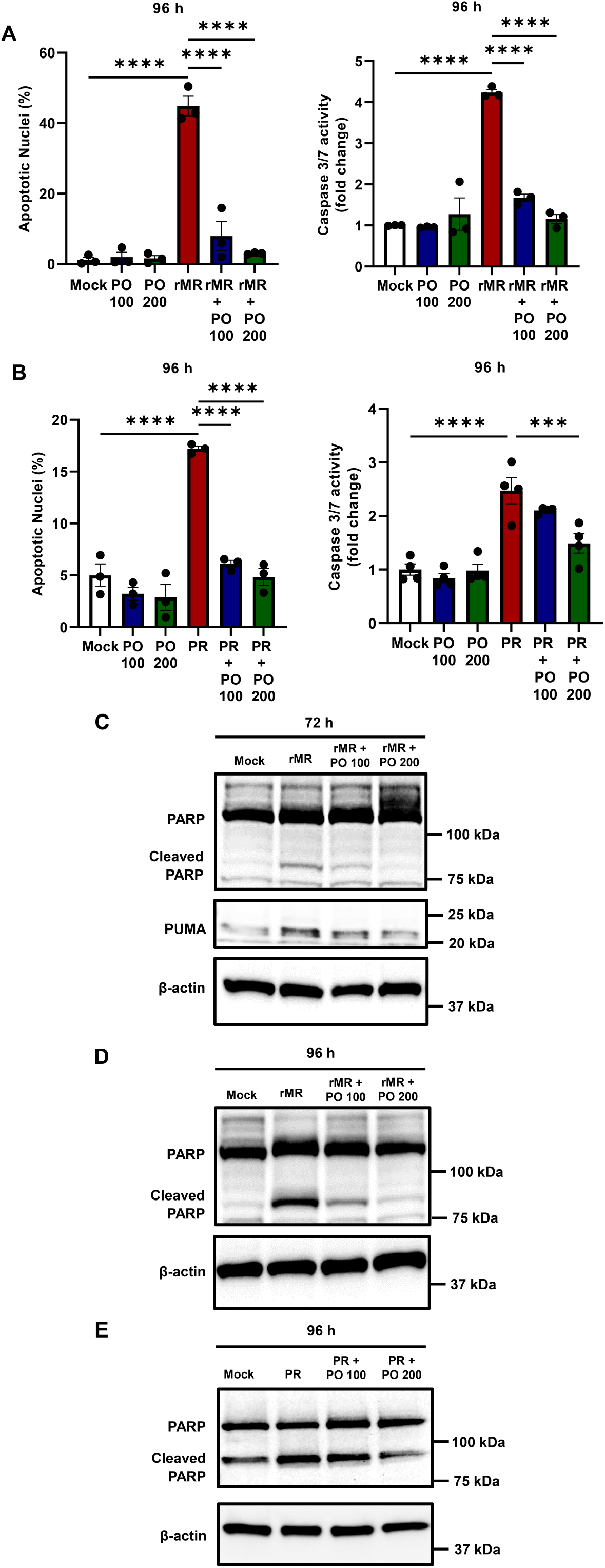
Palmitoleate (PO) protects against ZIKV-induced apoptosis in neuronal cells. **A, B.** SH-SY5Y cells infected with 1.0 MOI of rMR showed increased percent apoptotic nuclei and caspase 3/7 activity compared to mock-infected cells. Treatment of PO (100 µM and 200 µM) significantly protects against the rMR-induced increase in percent apoptotic nuclei and caspase 3/7 activity due at 96 h of post-infection(hpi). **C.** Immunoblot analysis with the treatment of PO showed decreased cleaved PARP and PUMA expression at 72 h when infected with 0.1 MOI rMR. **D.** Further, cleaved PARP was also decreased with the PO treatment at 96 h with rMR infection. **E.** Treatment of 200 µM PO also decreased the levels of cleaved PARP that was increased with 0.1 MOI PR strain infection in SH-SY5Y cells. Beta-actin was used as loading control and remained unchanged. The images are representative images. Data presented as mean ± SEM, n=3 or 4 *** p<0.001, **** p<0.0001 compared to mock or ZIKV infected cells.

### Palmitoleate mitigates ZIKV-induced ER stress

#### PO mitigates rMR strain-induced ER stress

To determine the mechanism of PO protection against ZIKV infection, we infected neuronal cells with 1 MOI rMR and treated them without and with PO (100 200 µM) and assessed the mRNA levels of ER stress mediators. ER stress mediators such as s-XBP1, unspliced (us)-XBP1, CHOP, Bip, and ATF4 were evaluated after different time points for ZIKV infection (24 - 96 h). We observed an upregulation of s-XBP1 mRNA levels with 1 MOI rMR infection, whereas treatment of PO (100 µM and 200 µM) showed a significant reduction in the levels of s-XBP1 at 96 h post-infection (**Fig. 4A**). A similar non-significant trend of decreased s-XBP1 with PO treatment (100 µM and 200 µM) was observed at 24 h (**Fig. S1 A, B**). rMR infection showed a trend towards a decrease in us-XBP1 mRNA expression compared to the mock-infected cells (**Fig. 4B**). However, when normalized with total (t)-XBP1, rMR infection showed a significant decrease in the mRNA levels of us-XBP1 (**Fig. S2**). Treatment of PO to rMR-infected neuronal cells showed an increase in the trend of un-XBP1 mRNA expression compared to rMR-infected neuronal cells (**Fig. S2**).

**Fig. 4.**
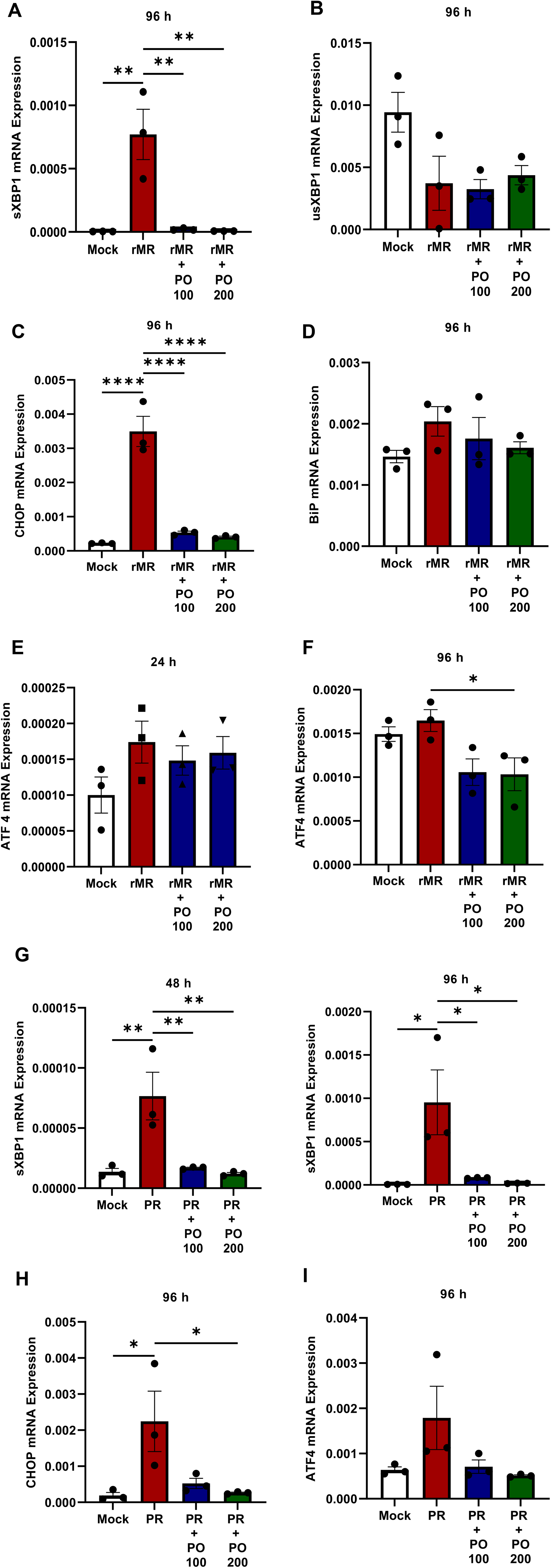
Palmitoleate mitigates ZIKV-induced ER stress. SH-SY5Y cells were infected with 1 MOI rMR or PR and treated with PO (100 µM and 200 µM) for different time points (24 h – 96 h). **A.** Cells infected with rMR significantly increased the mRNA expression of s-XBP1 compared to the mock and PO treatment drastically decreased the expression at 96 h. **B.** An insignificant decrease in us-XBP1 was observed with rMR infection at 96 h**. C.** rMR infection significantly increased the CHOP mRNA expression compared to mock but the PO treatment decreased the expression at 96 h. **D.** rMR infection led to an insignificant increase in Bip and PO treatment insignificantly decreased this expression at 96 h. **E.** rMR infection insignificantly increased ATF4 mRNA expression at 24 h and **F.** 200 µM of PO treatment significantly decreased the ATF4 expression at 96 h. **G.** Cells infected with PR drastically increased the s-XBP1 mRNA expression compared to the mock and PO treatment significantly decreased the expression at both 48 h and 96 h. **H.** CHOP mRNA expression was increased with PR infection and decreased with the PO treatment at 96 h. **I.** ATF4 expression was insignificantly increased with PR infection and PO treatment showed a decrease in trend at 96 h; relative to 18S rRNA. Data presented as mean ± SEM, n=3 *p<0.05, ** p<0.01, **** p<0.0001 compared to mock or ZIKV infected cells.

During sustained ER stress, s-XBP1 translocate to the nucleus and can increase the transcription of CHOP, a pro-apoptotic transcription factor. We studied the protective role of PO against CHOP expression in rMR-infected cells with and without the treatment of PO (100 - 200 µM). rMR-infected neuronal cells showed significantly increased CHOP mRNA expression after 96 h of post-infection compared to mock cells; however, treatment of PO, 100 - 200 µM significantly decreased the levels of CHOP mRNA (**Fig. 4C**). We also observed a noticeable trend towards an increase in CHOP mRNA expression with rMR infection and treatment of PO showed a trend towards decrease after 24 h of post-infection (**Fig. S3**). Bip mRNA expression, after 24 h, did not show any difference with the rMR infection or with the treatment of PO (**Fig. S4**). However, after 96 h a pattern of increase in Bip mRNA expression was observed in the rMR-infected group, and treatment of PO (100 µM and 200 µM) showed an insignificant decrease (**Fig. 4D**). A similar pattern was observed for ATF4 mRNA expression with rMR infection and PO treatment after 24 h of post-infection time point (**Fig. 4E**). However, after 96 h post-infection, 200 µM PO treatment significantly decreased the mRNA expression of ATF4 compared to rMR-infected neuronal cells (**Fig. 4F**).

#### PO mitigates PR strain-induced ER stress

Neuronal cells infected with 1 MOI of PR strain also showed a significant increase in s-XBP1 mRNA expression (**Fig. 4G**). Remarkably, treatment of 100 – 200 µM PO significantly decreased the expressions of s-XBP1 after both 48 h and 96 h post-infection time points (**Fig. 4G**). Further, PR infection increased the levels of CHOP mRNA, and treatment of PO significantly decreased the CHOP expression in neuronal cells (**Fig. 4H**). A trend towards an increase in ATF4 mRNA expression was noted with PR infection, whereas PO treatment exhibited an insignificant decrease in the ATF4 mRNA levels at 96 hpi (**Fig. 4I**). Together, these data suggest that treatment of PO mitigated both rMR and PR strain-induced activation of ER stress.

### Palmitoleate decreases protein markers of ER stress with ZIKV infection

To further determine whether PO can also mitigate the protein expression of ER stress mediators, we infected neuronal cells with 0.1 MOI rMR and treated them with PO (100 µM and 200 µM) at 8 h, 72 h, and 96 h of post-infection time points (**Fig. 5A-E**). We observed increased phosphorylated forms of IRE1α (p-IRE1α) activation with 0.1 MOI rMR infection at 8 h of post-infection (**Fig. 5A**). Treatment with 200 µM PO resulted in decreased levels of p-IRE1α (**Fig. 5A**). However, levels of total (t)-IRE1α did not change in all the treatment conditions. Similarly, an increased level of cleaved ATF6 was observed at 8 h with rMR infection, and treatment of 200 µM PO exhibited a decrease in cleaved ATF6 levels (**Fig. 5B**).

**Fig. 5.**
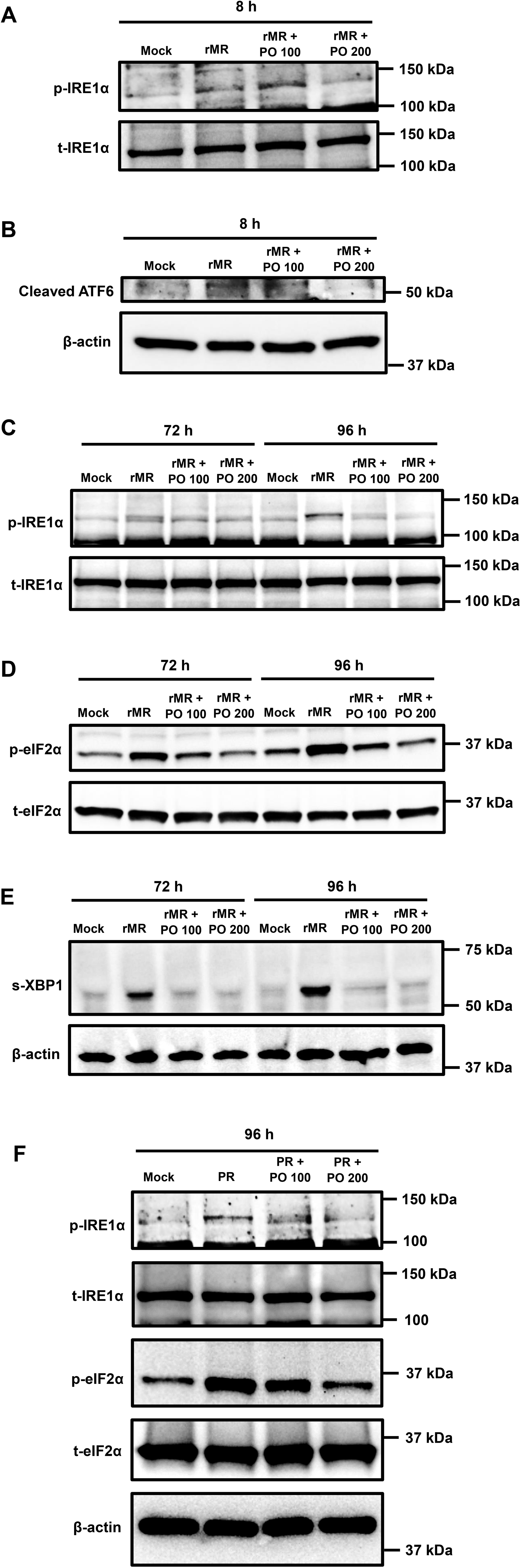
Palmitoleate decreases protein markers of ER stress with ZIKV infection. SH-SY5Y cells were infected with 0.1 MOI rMR or PR and treated with PO (100 µM and 200 µM) at different post-infection time points (8 h, 72 h, 96 h). **A.** Immunoblot analysis showed that PO treatment of 200 µM decreased the p-IRE1α. **B.** cleaved ATF6 at 8 h of post-infection time point when infected with rMR. PO treatment (100 µM and 200 µM) decreased the **C.** p-IRE1α, **D.** p-eIF2α, and **E.** s-XBP1 at both 72 h and 96 h of post-infection time point when infected with rMR. **F.** PO treatment (100 µM and 200 µM) decreased the p-IRE1α and p-eIF2α when infected with PR at 96 h of post-infection time point.

Alarmingly, we also observed substantial increases in p-IRE1α after 72 h and 96 h of post-infection time points, but treatment of PO (100-200 µM) drastically decreased the levels of p-IRE1α (**Fig. 5C**). Further, increased levels of p-eIF2α and s-XBP1 were observed with rMR infection at both 72 h and 96 h of post-infection time points (**Fig. 5D, E**). However, treatment of PO (100 µM and 200 µM) dramatically decreased the expressions of p-eIF2α and s-XBP1 levels (**Fig. 5D, E**).

Similar to rMR strain infection, neuronal cells infected with 0.1 MOI PR strain of ZIKV also showed a drastic increase in the levels of p-IRE1α and p-eIF2α. Treatment of PO (100 µM and 200 µM) decreased the expression of p-IRE1α and p-eIF2α at 96 h post-infection (**Fig. 5F**). However, the levels of t-IRE1α, t-eIF2α, and β-actin levels remained unchanged in all the treatment conditions (**Fig. 5 F**). In summary, these results suggest that treatment of PO prevents the activation of sustained ER stress with both rMR and PR strain infection to neuronal cells.

### Palmitoleate attenuates viral envelope levels and infectious ZIKV titer in neuronal cells

To further determine the protective role of PO against ZIKV infection, we infected neuronal cells with 1 MOI rMR, and at 96 h of post-infection time point, we observed a significant increase in the Zika viral envelope (E) RNA copy number (**Fig. 6A**). However, treatment of PO (100 µM and 200 µM) significantly decreased the levels of viral *E gene* RNA copy number (**Fig. 6A**). Immunoblot analysis of ZIKV-infected neuronal cells with 0.1 MOI rMR resulted in a considerable increase in viral envelope protein expression after 48 h and 72 h of post-infection time points. However, treatment of PO (100 µM and 200 µM) attenuated the levels of the viral envelope in neuronal cells (**Fig. 6B**). Similarly, 0.1 MOI PR infection increased the expression of ZIKV envelope protein at 96 h and treatment of PO (100 µM and 200 µM) markedly decreased the envelope protein expression (**Fig. 6C**). Further, immunofluorescence analysis of Zika viral E protein showed an increase in ZIKV envelope protein levels in neuronal cells infected with 1 MOI rMR compared to the mock-infected cells, but 200 µM of PO markedly ameliorated the viral envelope protein levels at 48 hpi time point (**Fig. 6D**). Plaque assay was employed to determine the infectious titer with ZIKV infection in neuronal cells. Cell culture supernatant collected from 0.1 MOI PR infected cells showed a significant increase in plaque forming units (PFUs) and 200 µM PO treatment significantly decreased the PFUs (**Fig. 6E**). Similarly, cells infected with 1 MOI rMR showed drastically increased PFUs at 96 h, whereas treatment of PO (100 µM and 200 µM) significantly decreased the infectious virus titer (**Fig. 6F**).

**Fig. 6.**
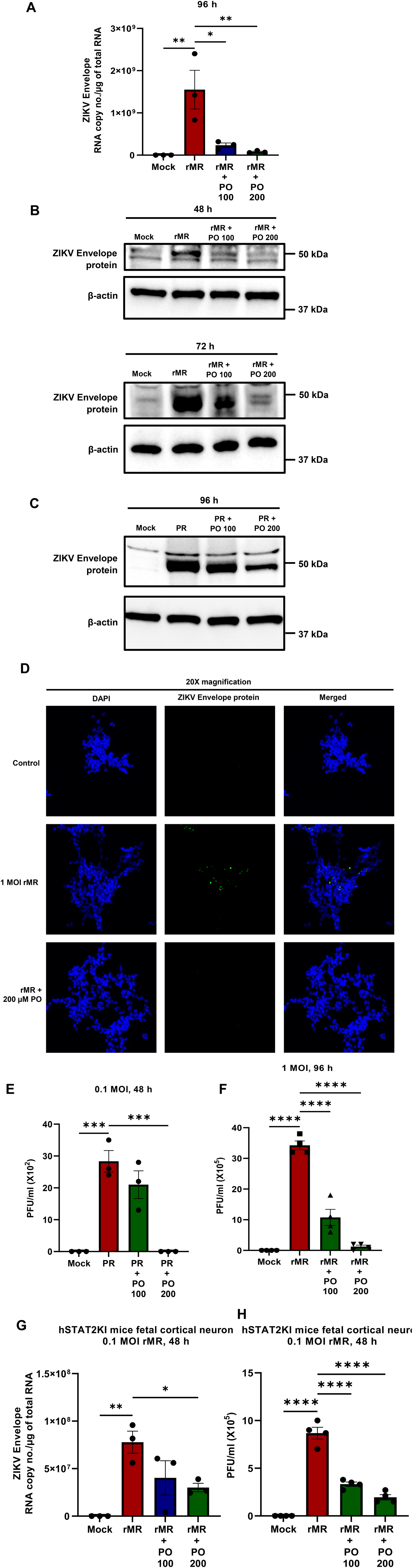
Palmitoleate attenuates viral envelope levels and infectious ZIKV titer in neuronal cells. **A.** SH-SY5Y cells infected with 1 MOI rMR showed a drastic increase in the Zika viral envelope RNA copy number compared to the mock and treatment of palmitoleate (PO, 100 µM and 200 µM) drastically decreased the viral RNA copy number. **B.** Immunoblot analysis showed SH-SY5Y cells infected with 0.1 MOI rMR expressed high ZIKV envelope protein at both 48 h and 72 h of post-infection time points and PO treatment (100 µM and 200 µM) attenuated the viral protein expression. **C.** 0.1 MOI PR infection increased the envelope protein expression after 96 h of post-infection time point, whereas PO treatment decreased this expression. **D.** Immunofluorescence analysis showed that 1 MOI rMR infection in SH-SY5Y cells increased viral envelope protein staining and 200 µM PO treatment decreased the staining at 48 h. Beta-actin was used as loading control and remained unchanged. The images are representative images. **E.** Plaque assay showed cell culture supernatant of SH-SY5Y cells infected with 0.1 MOI PR strain resulted in a significant increase in Plaque Forming Units (PFUs) and 200 µM PO treatment decreased the PFU at 96 h. **F.** PO treatment (100µM and 200µM) significantly attenuated the PFUs when the cells were infected with 1 MOI rMR at 96 h. **G.** 0.1 MOI rMR infection in primary cortical neurons from hSTAT2KI mice fetus showed dramatically increased ZIKV envelope RNA copy number relative to total cellular RNA and treatment with 200 µM PO significantly decreased the viral copy number. **F.** Plaque assay showed rMR infection increased the PFU and treatment of PO (100 µM and 200 µM) decreased the PFUs in primary cortical neurons isolated from hSTAT2KI mice. Data presented as mean ± SEM, n=3 or 4, *p<0.05, ** p<0.01, *** p<0.001, **** p<0.0001 compared to mock or ZIKV infected cells.

Further to test the role of PO in attenuating ZIKV replication in primary cells we isolated cortical neurons from hSTAT2KI mice fetuses and infected them with 0.1 MOI rMR. At 48 h of post-infection, a significant increase in ZIKV envelope RNA copy number was detected in the supernatant of the infected cells, whereas treatment with 200 µM of PO significantly decreased its expression (**Fig. 6G**). Further, plaque assay showed that rMR infection has significantly increased the PFUs, whereas treatment of PO (100 µM and 200 µM) drastically decreased the ZIKV infectious titer in hSTAT2KI fetal cortical neurons culture supernatant (**Fig. 6H**). These data suggest that treatment of PO attenuates ZIKV envelope RNA and protein expression; further, decreasing the infectious viral titer in both neuronal cells and primary cortical neurons isolated from hSTAT2KI mice fetuses.

## DISCUSSION

In this study, we have identified the activation of ER stress and apoptosis with ZIKV infection in neuronal cells and established the protective role of palmitoleate, an omega-7 monounsaturated fatty acid against ZIKV-induced ER stress and apoptosis in neuronal cells, *in vitro*. The principle findings of this manuscript are illustrated in Fig. 7, which are 1) ZIKV infection induces caspase-dependent apoptosis, 2) ZIKV infection upregulates pro-apoptotic markers such as Bim, and PUMA, and decreases the expression of anti-apoptotic markers namely Bcl-2, Bcl-xL, and Mcl-1 and can cause activation of the intrinsic pathway of apoptosis, 3) ZIKV infection also induces the activation of ER stress, 4) Treatment of palmitoleate protects against ZIKV-induced ER stress and apoptosis, and 5) Palmitoleate attenuates viral replication and its release in both neuronal cells and primary cortical neurons isolated from hSTAT2KI mice fetus.

**Fig. 7.**
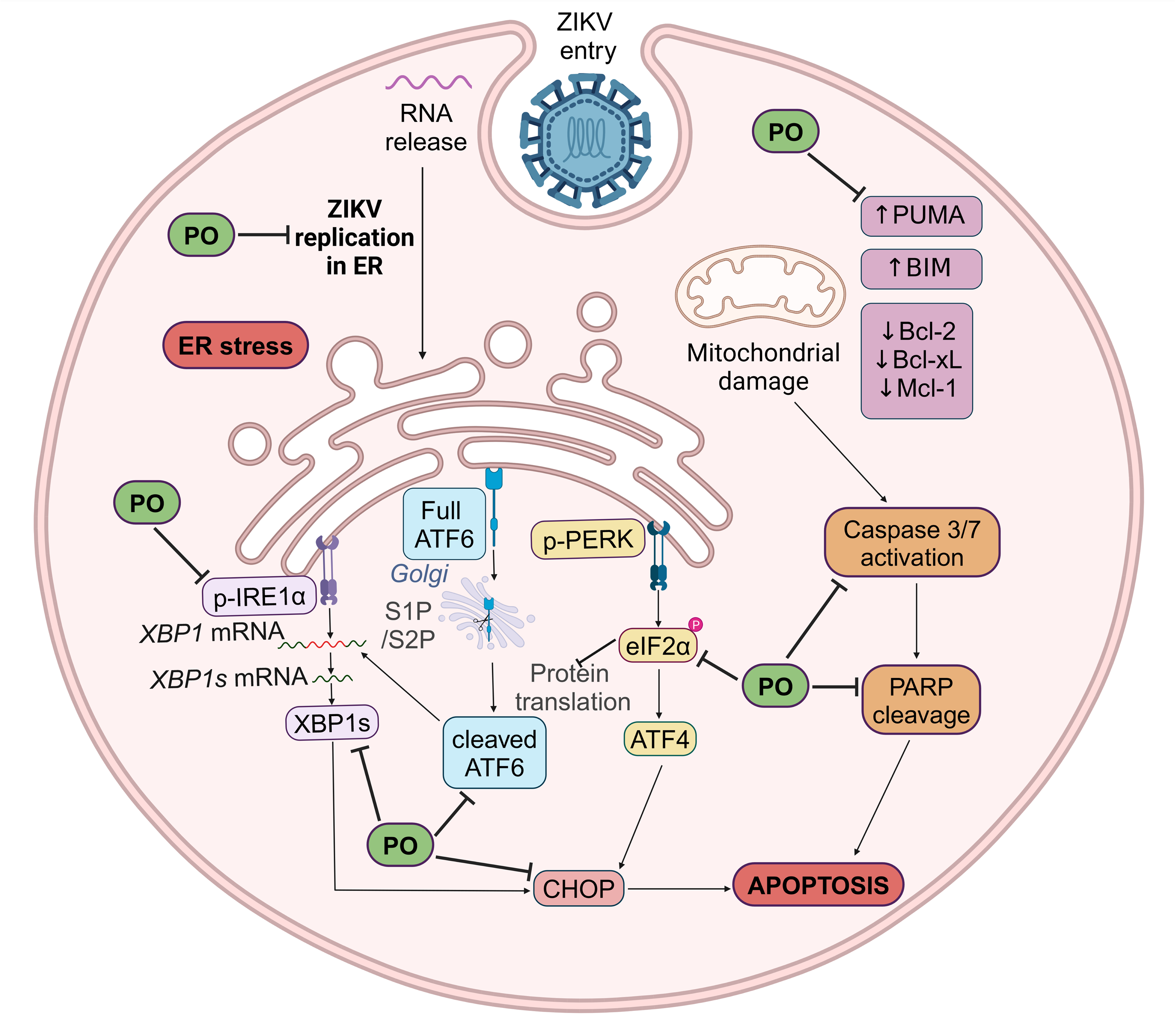
Schematic representation of ZIKV-induced ER stress and apoptosis in neuronal cells and the protective role of palmitoleate. ZIKV infection induces caspase-dependent apoptosis and sustained ER stress activation in the neuronal cells. An increase in mitochondrial pro-apoptotic mediators BIM, and PUMA, and a decrease in anti-apoptotic mediators Bcl-2, Bcl-xL, and Mcl-1 are observed due to the ZIKV infection. Further, ZIKV infection results in the activation of caspases and increases the levels of cleaved PARP leading to neuronal apoptosis. Infection with ZIKV also causes sustained ER stress as evidenced by an increases the phosphorylation of IRE1α and eIF2α, cleavage of ATF6, splicing of XBP1, and CHOP. In contrast, when the ZIKV-infected neuronal cells are treated with palmitoleate it results in dramatic decrease in PUMA, caspase 3/7 activation, and cleaved PARP expressions. Similarly, the treatment of palmitoleate also reduces ER stress which is evidenced by a decrease in p-IRE1 α, p-eIF2 α, s-XBP1, cleaved ATF6, and CHOP. Further, treatment of palmitoleate also decreases the ZIKV replication in the neurons.

During pregnancy, the placental and blood-brain barrier protects the fetal brain from pathogens and the ZIKV breaches these barriers to get into the fetal brain(20). Our previous study showed that ZIKV infections in placental trophoblasts resulted in apoptosis by way of activation of cellular stress pathways(12). ZIKV infection in the chorionic villi dissected from term human placental explants showed an increase in ZIKV RNA at 12 h and peaks at 72 h of post-infection and resulted in placental villi apoptosis(21). In the present study, we observed that ZIKV induces apoptosis in neuronal cells, and it causes caspase-dependent apoptosis. During ZIKV infection, there are several mechanisms involved in neuropathogenesis; neuronal apoptosis is the major cell death type observed during the developmental brain disorder(20). ZIKV infection induces cell death in human neural progenitor cells which was evidenced by an increase in caspase 3 activation; further bulk RNA-sequencing showed upregulation of pro-apoptotic genes(22). ZIKV infection in neural stem cells also causes a caspase 3/7 mediated-cell death(23). Further, ZIKV infection in mouse embryonic brain at 13.5 embryonic days resulted in the activation of caspase 3 in cells located in the cortical plate and intermediate zone(24). Gene ontology analysis of the whole brain from ZIKV-infected littermate showed upregulation of genes involved in apoptotic pathways(24). Further, offspring from ZIKV-infected pregnant mice showed intrauterine growth restriction phenotype and upregulation of genes related to apoptosis and autophagy like Casp6, Bcl2, Huntingtin, Bcl2 modifying factor, ABL proto-oncogene 1 (Abl1), and Immunity-related GTPase family M member 1 (Irgm1)(25). Fetal brain tissues with microcephaly due to the ZIKV infection had increased caspase 3 levels in the parenchyma which was contained in the cerebral cortex region(26). Neuroendocrine PC12 cells overexpressed with ZIKV envelope protein also showed an increase in cleaved caspase 3 and 9 protein expression which resulted in the activation of the intrinsic pathway of apoptosis (27). These previous results are consistent with the present study as rMR and PR infections to neuronal cells result in an increased caspase activity and induce neuronal apoptosis.

In the present study, we also analyzed the expression pattern of the pro-apoptotic BH3-only domain-containing protein, Bim, and it was compared to the anti-apoptotic Bcl2 family of proteins such as Bcl-2, Bcl-xl, and Mcl-1 from 8 h to 72 h of post infections. Since Bax and Bak can be activated by BH3-only proteins such as Bim; the increased Bim & PUMA and decreased anti-apoptotic expressions with ZIKV infection in neuronal cells is similar to previous studies which showed the Bax and Bak activation along with decreased anti-apoptotic expression with ZIKV infection(27, 28). Earlier studies have focused on the correlation between Bax and Bak with anti-apoptotic markers(27, 28). ZIKV infection causes activation of Bax and its oligomerization in the mitochondria(11). Further, Bax knockdown in neuronal cells drastically decreased apoptosis and neuronal cells overexpressed with Flag-Bcl-xL had greatly decreased the Bax activation(11). During *Flavivirus* infection treatment with the small molecule inhibitor for Bcl-xL resulted in enhanced Bax/Bak-dependent apoptosis and attenuated the expression of Mcl-1(28). Further, overexpression of ZIKV envelope protein in neuroendocrine PC12 cells led to decreased levels of Bcl-2 and increased Bax expression for the initiation of apoptosis(27). Together our confirms that during ZIKV infection to neuronal cells showed increased expression of mitochondrial pro-apoptotic mediators and decreased levels of anti-apoptotic mediators.

We have also demonstrated that ZIKV infection induces ER stress in neuronal cells. CHOP is an important pro-apoptotic marker for sustained ER stress-mediated apoptosis. In the present study, we observed increased CHOP expression with the ZIKV infection. Previous studies have shown that CHOP activation during ER stress can induce upregulation of pro-apoptotic BH3 mitochondrial mediators such as PUMA(29, 30). However, further studies are needed to test the mechanism and the role of CHOP in the mitochondrial intrinsic pathway of apoptosis in ZIKV-infected neuronal cells.

Human occipital cortex from ZIKV-infected fetuses with microcephaly and human neural stem cells infected with ZIKV showed an increase in ER stress and UPR(31). ZIKV infection in SK-N-SH cells (parental neuronal cell of SH-SY5Y) showed activation of p-IRE1α, s-XBP1, p-eIF2α, cleaved ATF6 and CHOP; further AG6 *Ifnagr^−/−^* mice infected with ZIKV showed activation of ER stress as evidenced by an increase in p-IRE1α, s-XBP1, cleaved ATF6, ATF4, and CHOP(32). The outcomes observed in earlier studies are in line with the present findings proving that ZIKV infection in neuronal cells resulted in activation of p-IRE1α, s-XBP1, p-eIF2α, cleaved ATF6, and CHOP. Further, we showed that inhibition of eIF2α and IRE1α blocked ZIKV infection-induced apoptosis which proved the critical role in activation of ER stress during neuronal apoptosis. Collectively, our studies showed mechanistic evidence and a critical role in the activation of ER stress in ZIKV-induced apoptosis.

In the present study, ZIKV infection in neuronal cells resulted in increased cleaved ATF6 which could be due to the translocation of ATF6 from ER to Golgi followed by the cleavage of ATF6 by S1P/S2P proteases(33). Studies have shown that the IRE1α/XBP1 signaling can be enhanced independently by PERK/ATF4 and ATF6 arms of ER stress suggesting a crosstalk between different arms of ER stress(34, 35). Thus, during ZIKV infection cleaved ATF6 might increase XBP1 mRNA for its further splicing. Studies have also shown that p-eIF2α results in the translation of ATF4(36) and that PERK/ATF4-dependent increases the expression of IRE1α which further improves the XBP1 splicing efficiency(35). In the present study, we observed increased phosphorylation of eIF2α due to the ZIKV infection which might have resulted due to increased IRE1α expression and can be a contributing factor for the increased expression of s-XBP1 mRNA and protein.

ZIKV induces caspase-dependent and ER stress-related apoptosis in neuronal cells. Although IRE1α endonuclease and eIF2α dephosphorylation inhibitors significantly attenuate ZIKV-induced apoptosis, they could have potential cytotoxicity in the developing embryo. Therefore, our goal is to use the dietary nutrient compound palmitoleate (PO) to mitigate ZIKV infection-induced ER stress and apoptosis. PO decreased the palmitate-induced hepatocyte lipoapoptosis and attenuated the PUMA and Bim expressions(37). Our previous study showed that PO treatment decreased the ZIKV infection and protected ZIKV-infected trophoblasts from apoptosis and ER stress by decreasing the downstream targets of ER stress markers such as CHOP and s-XBP1(12). Likewise, in the present study, we observed the protective effects of PO in decreasing ZIKV-induced apoptosis and ER stress in the neuronal cells against two ZIKV strains namely, rMR and PR.

An increase in viral protein synthesis during infection in a host cell can affect the folding capacity of the ER which results in the activation of ER stress(38). Previous studies have shown that biochemical compounds involved in attenuating ZIKV replication of which isoquercitrin treatment prevents the ZIKV internalization into the host cell and naringenin could prevent viral assembly or replication(16, 39, 40). In the present study, we observed the treatment of PO dramatically mitigated ZIKV replication and infectious ZIKV titer in neuronal cells and cortical neuronal cells from hSTAT2KI mice fetuses. Further studies are required to identify the molecular mediators of palmitoleate in attenuating ZIKV replication, ER stress, and apoptosis.

Earlier studies have shown that ZIKV infection altered lipid metabolism in the host cell(41, 42) and supplementation of unsaturated fatty acids like oleic acid induces an increase in lipid droplets (LD) (43). Firstly, ZIKV infection in human neuronal cells resulted in increased LD accumulation due to a decrease in the enzymes involved in triglyceride lipolysis like hormone-sensitive lipase (HSL), and adipose triglyceride lipase (ATGL)(41). Additionally, an increase in lipogenic proteins like acyl-CoA: diacylglycerol acyltransferase-1 (DGAT-1), fatty acid synthase (FASN), peroxisome proliferator-activated receptor-γ (PPAR-γ), sterol regulatory element-binding protein 1 (SREBP-1), and perilipin 2 (PLIN-2) were also observed(41). Further, a decrease in viral replication was evidenced when LD formation was blocked by targeting DGAT-1 inhibition in both *in vitro* and *in vivo* brain sections obtained from ZIKV-infected animals(41). These observations suggest that ZIKV-induced LD biogenesis is critical for viral replication and pathogenesis in neuronal cells(41). ZIKV infection after 24 h of infection showed an increased number and size of LDs in less differentiated human neuronal progenitor cells but no significant changes in LD were observed at later points such as 48 h and 56 hpi(42). Similarly, poly I:C, which mimics RNA virus infection also resulted in an increased accumulation of LDs in the astrocytes in early time points, but after 72 h of treatment a drastic decrease in LDs number was observed(43). Overall, these studies show that an increase in LD accumulation was observed during the initial stages of ZIKV infection, and at the later time points it decreased. Secondly, immortalized primary astrocytes pre-treated with oleic acid before ZIKV infection resulted in increased lipid droplet (LD) accumulation(43). These LDs due to the treatment of oleic acid subsequently induced the interferon response, when exposed to ZIKV infection and resulted in a decreased ZIKV mRNA expression(43). Likewise, in the present study, we speculate that treatment of PO could have increased the lipid droplet accumulation in the cells that might trigger the interferon responses resulting in decreased ZIKV replication, apoptosis, and ER stress in the neuronal cells. However, future studies are required to investigate the role of PO in the activation of interferon response and whether supplementation of PO could alter LD accumulation in the neuronal cells.

In conclusion, our mechanistic data suggests that ZIKV infection of neuronal cells results in the activation of sustained ER stress, activation of mitochondrial intrinsic pathway of apoptosis, and caspase-dependent apoptosis. Interestingly, treatment of PO significantly attenuated the activation of all three arms of ER stress and their downstream targets. Treatment of PO decreases the viral replication, and this could have led to less viral protein accumulation in ER thereby decreasing the ER-stress-mediated apoptosis. Further, treatment with PO also attenuated the PUMA expression proving PO’s role in preventing intrinsic pathway-mediated apoptosis. However, the exact mechanism of palmitoleate protection against ZIKV infection requires further investigation. Further, to translate the protective role of PO into clinical practice, it is critical to study its effect against ZIKV infection *in vivo* with animal models and subsequently in cohort studies with at risk populations of pregnant women.

## List of abbreviations

ZIKV: Zika virus
rMR: recombinant clone of African strain, MR766
PR: PRVABC59 Asian strain
MOI: multiplicity of infection
PARP: poly (ADP-ribose) polymerase
MCL-1: Myeloid cell leukemia 1
BCL-2: B-cell lymphoma 2
Bcl-xL: B-cell lymphoma-extra large
ATF4: Activating transcription factor 4
ATF6: Activating transcription factor 6
BIM: BCL2-interacting mediator of cell death
CHOP: C/EBP homologous protein
DAPI: 4’-6-diamidino-2-phenylindole dihydrochloride
ER: Endopasmic reticulum
PO: Palmitoleate
eIF2α: Eukaryotic initiation factor α
IRE1α: Inositol-requiring transmembrane kinase/endoribonuclease 1α
MAPK: Mitogen activated protein kinase
PERK: protein kinase RNA-like endoplasmic reticulum kinase
XBP1: X-box binding protein 1
Bip: immunoglobulin heavy chain binding protein
PUMA: p53 upregulated modulator of apoptosis
JNK: c-Jun N-terminal kinase
Bax: Bcl-2-associated X protein
Bak: Bcl-2 antagonist killer 1
hSTAT2KI: humanized Signal Transducer and Activator of Transcription 2 knockin
PFU: plaque-forming unit
LD: lipid droplets
SEM: standard error of the mean

## Data Availability

All the data generated or analyzed during this study are included in the article main text and in supplementary information files.

## Acknowledgments

We would like to thank Terri Fangman in the microscope core at the Nebraska Center for Biotechnology, University of Nebraska-Lincoln.

