## Supplementary files for "Palmitoleate Protects against Zika virus infection-induced Endoplasmic Reticulum Stress and Apoptosis in Neurons"

[1]Department of Nutrition and Health Sciences, University of Nebraska-Lincoln, Lincoln, NE, USA; [2]Graduate Interdisciplinary Program Neuroscience, University of Arizona, Tucson, AZ, USA; [3]Department of Pediatrics, University of Nebraska Medical Center, Omaha, NE, USA; [4]College of Allied Health Professions Medical Nutrition Education, University of Nebraska Medical Center, Omaha, NE, USA; [5]Department of Biochemistry, University of Nebraska-Lincoln, Lincoln, NE, USA.

*Address for Correspondence: Sathish Kumar Natarajan, PhD
Associate Professor,
Department of Nutrition & Health Sciences
University of Nebraska-Lincoln
229 Filley Hall, Lincoln, NE 68583-0806

ORCID: orcid.org/0000-0001-7491-8592

**Running title: Palmitoleate prevents ZIKV infection in neuronal cells.**

**Keywords: Congenital Zika Syndrome, Microcephaly, Nutrient Intervention, ER stress**

#### Figure S1

A

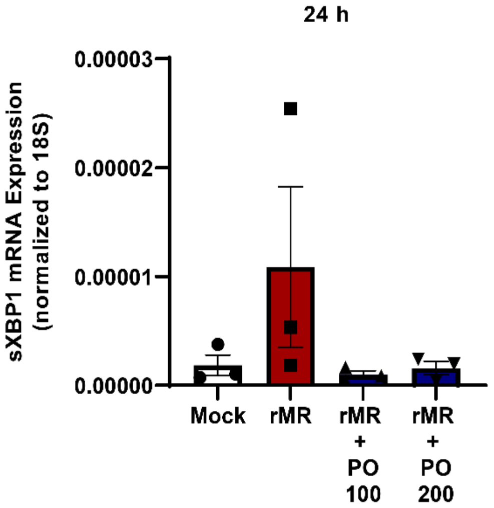


B

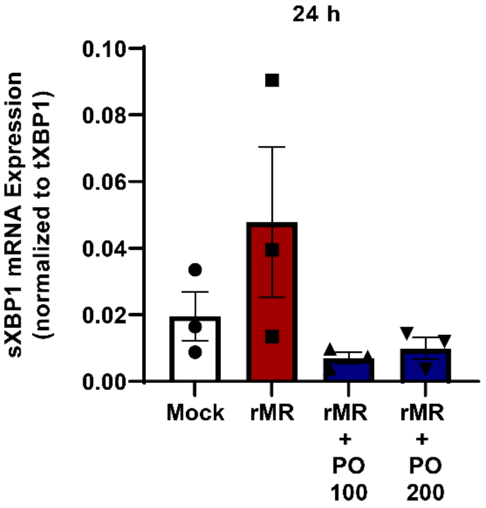


**Figure S1. s-XBP1 mRNA expression insignificantly increased with rMR infection and PO treatment trivially decreased the expression.** SH-SY5Y cells infected with 1 MOI rMR and treated with PO (100 – 200 µM) **A.** s-XBP1 mRNA levels were normalized to 18S and **B.** s-XBP1 mRNA levels were normalized to t-XBP1 at 24 h of post-infection time point. Data presented as mean ± SEM, n=3.

### Figure S2

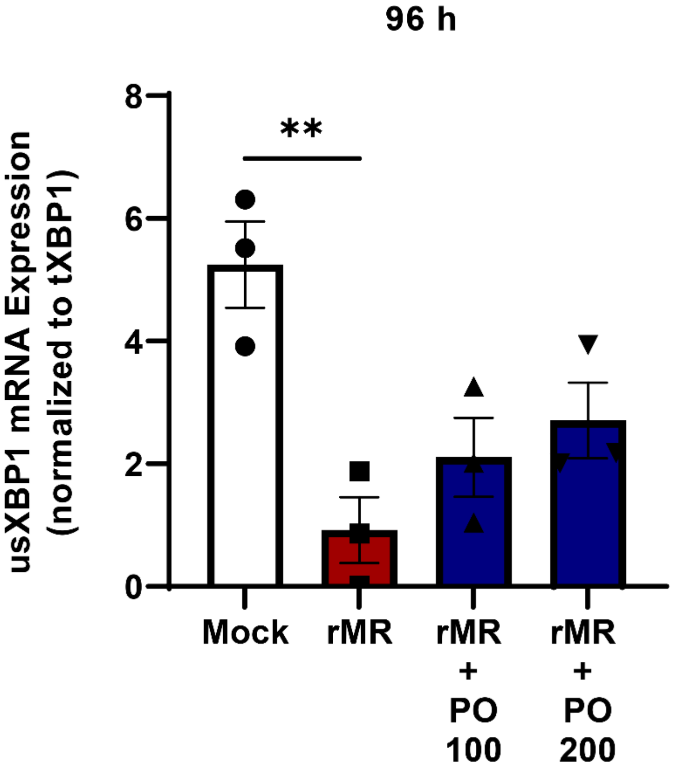


**Figure S2. Unspliced (us)-XBP1 mRNA expression significantly decreased with rMR infection and PO treatment insignificantly increased the expression.** SH-SY5Y cells infected with 1 MOI rMR and treated with PO (100 – 200 µM) at 96 h of post-infection time point showed a significant decrease in unspliced-XPB1 mRNA relative to total (t)XBP1. Data presented as mean ± SEM, n=3, ** $p<0.01$ compared to mock.

### Figure S3

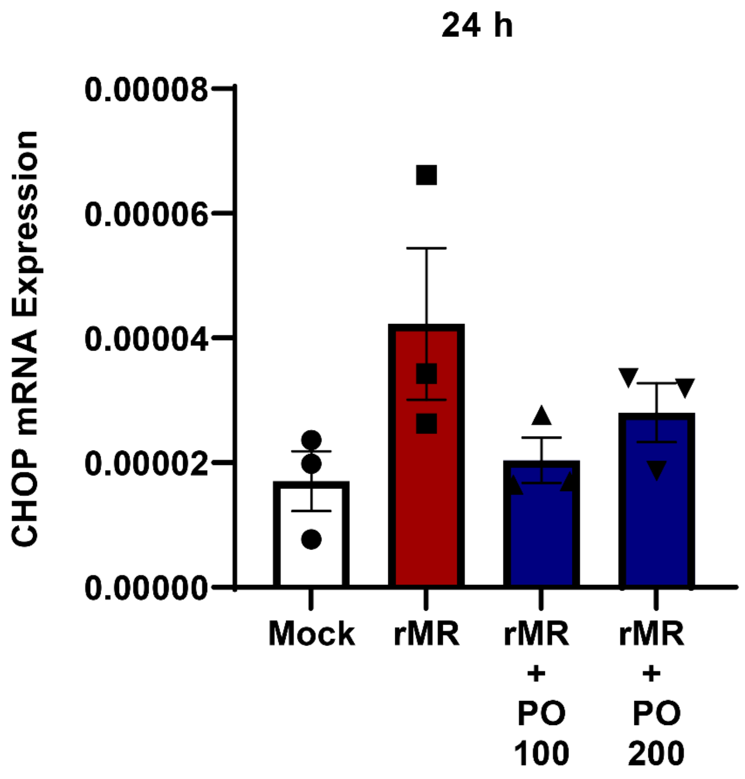


**Figure S3. CHOP mRNA expression showed an insignificant pattern of increase with rMR infection and treatment of PO slightly decreased the expression.** SH-SY5Y cells infected with 1 MOI rMR showed a non-significant trend towards an increase in CHOP mRNA levels relative to 18S rRNA and treatment of PO (100 – 200 µM) did not alter the mRNA levels of CHOP compared to mock infected cells at 96 h of post-infection time point. Data presented as mean ± SEM, n=3.

### Figure S4

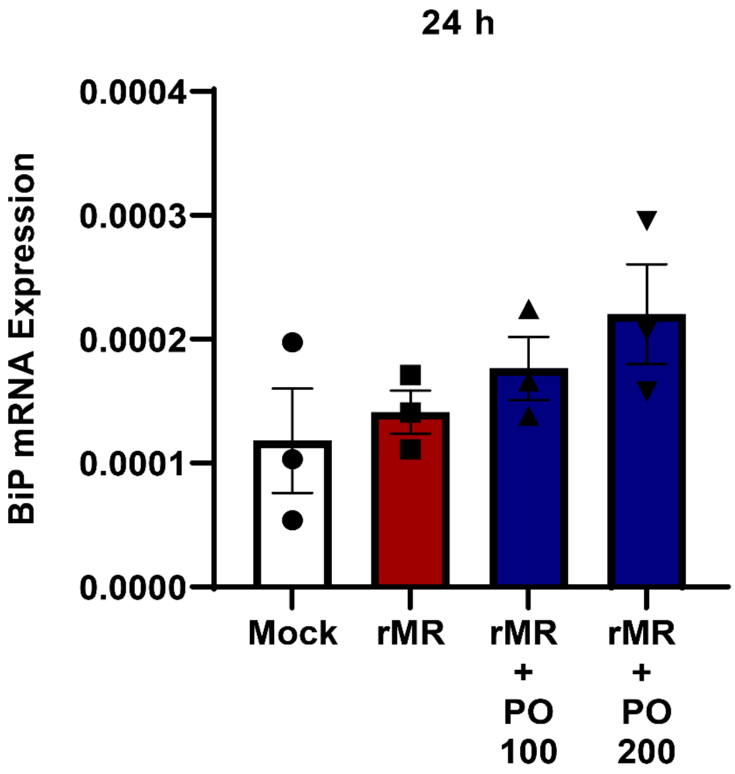


**Figure S4. Bip mRNA expression showed no difference in the pattern with rMR infection and PO treatment.** SH-SY5Y cells infected with 1 MOI rMR and treated with PO (100 – 200 µM) did not show any change at 24 h of post-infection time point. Data presented as mean ± SEM, n=3.
